# Rapid Brain Responses to Familiar vs. Unfamiliar Music – an EEG and Pupillometry study

**DOI:** 10.1101/466359

**Authors:** Robert Jagiello, Ulrich Pomper, Makoto Yoneya, Sijia Zhao, Maria Chait

## Abstract

Human listeners exhibit marked sensitivity to familiar music – perhaps most readily revealed by popular “name that tune” games, in which listeners often succeed in recognizing a familiar song based on extremely brief presentation. In this work we used electro-encephalography (EEG) and pupillometry to reveal the temporal signatures of the brain processes that allow differentiation between familiar and unfamiliar music. Participants (N=10) passively listened to snippets (750 ms) of familiar and, acoustically matched, unfamiliar songs, presented in random order. A group of control participants (N=12), which were unfamiliar with all of the songs, was also used. In the main group we reveal a rapid differentiation between snippets from familiar and unfamiliar songs: Pupil responses showed greater dilation rate to familiar music from 100-300 ms post stimulus onset. Brain responses measured with EEG showed a differentiation between familiar and unfamiliar music from 350 ms post onset but, notably, in the opposite direction to that seen with pupillometry: Unfamiliar snippets were associated with greater responses than familiar snippets. Possible underlying mechanisms are discussed.

## 1. Introduction

The human auditory system exhibits a marked sensitivity to, and memory of music (Fujioka, Trainor, Ross, Kakigi, & Pantev, 2005; Trainor, Marie, Bruce, & Bidelman, 2014; Filipic, Tillmann, & Bigand, 2010; Halpern & Bartlett, 2010; Schellenberg, Iverson, & McKinnon, 1999; Koelsch, 2018). The concept of music familiarity heavily relies on long term memory traces (Wagner, Shannon, Kahn, & Buckner, 2005) in the form of auditory mental imagery (Halpern & Zatorre, 1999; Kraemer, Macrae, Green, & Kelley, 2005; Martarelli, Mayer, & Mast, 2016) and is also linked to autobiographical memories (Janata, Tomic, & Rakowski, 2007). Our prowess towards recognizing familiar musical tracks is anecdotally exemplified by ‘Name That Tune’-games, in which listeners of a radio station are asked to name the title of a song on the basis of a very short excerpt. Even more fleeting recognition scenarios may occur when switching from one station to another while deciding which one to listen to—beloved songs often showing the ability to swiftly catch our attention, causing us to settle for a certain channel. Here we seek to quantify such recognition in a lab setting. We aim to understand how quickly listeners’ brains can identify familiar music snippets among unfamiliar snippets and pinpoint the neural signatures of this recognition. Beyond basic science, understanding the brain correlates of music recognition is useful for various music-based therapeutic interventions (Montinari, Giardina, Minelli, & Minelli, 2018). For instance, there is a growing interest in exploiting music to break through to dementia patients for whom memory for music appears well preserved despite an otherwise systemic failure of memory systems (Cuddy & Duffin, 2005; Hailstone, Omar, & Warren, 2009). Pinpointing the neural signatures of the processes which support music identification may provide a clue to understanding the basis of the above phenomena, and how it may be objectively quantified.

Previous work using behavioral gating paradigms demonstrates that the latency with which listeners can identify a familiar piece of music (pop or instrumental) amongst unfamiliar excerpts ranges from 100 ms (Schellenberg et al., 1999) to 500 ms (Bigand, et al, 2009; Filipic et al., 2010; Krumhansl, 2010; Tillmann et al, 2014). It is likely that such fast recognition is driven by our memory of the timbre and other spectral distinctivness of the familiar piece (Bigand et al, 2009; 2011; Agus et al, 2012; Suied et al, 2014).

According to bottom-up theories of recognition memory, an incoming stimulus is compared to stored information, and upon reaching a sufficient congruence is then classified as familiar (Bigand et al., 2009; Wixted, 2007). A particular marker in the EEG literature that is tied to such recognition processes is the late positive potential (LPP; Curran, 2000; Johnson, 1995; Rugg & Curran, 2007): The correct identification of a familiar stimulus typically results in a sustained positivity ranging from 500 to 800 ms post stimulation in left central-parietal regions, which is absent for unfamiliar stimuli (Kappenman & Luck, 2012). This parietal old versus new effect has consistently been found across various domains, such as facial (Bobes, Martín, Olivares, & Valdés-Sosa, 2000) and voice recognition (Zäske, Volberg, Kovács, & Schweinberger, 2014) as well as paradigms that employed visually presented- (Sanquist, Rohrbaugh, Syndulko, & Lindsley, 1980) and spoken words as stimuli (Klostermann, Kane, & Shimamura, 2008). In an fMRI study, Klostermann et al. (2009) used 2-second long excerpts of newly composed music and familiarized their participants with one half of the snippet sample while leaving the other half unknown. Subsequently, subjects were exposed to randomized trials of old and new snippets and were asked to make confidence estimates regarding their familiarity. Correct identification of previously experienced music was linked to increased activity in the posterior parietal cortex (PPC). However, due to the typically low temporal resolution of fMRI the precise time course of the recognition process remains unknown.

Pupillometry is also increasingly used as a measure of music recognition and, more generally, of the effect of music on arousal. This is part of a broader understanding that cognitive states associated with working memory load, vigilance, surprise or processing effort can be gleaned from measuring task-evoked changes in pupil diameter (Beatty, 1982; Bradley, Miccoli, Escrig, & Lang, 2008; Hess & Polt, 1964; Kahneman & Beatty, 1966; Mathôt, 2018; Preuschoff, Hart, & Einhäuser, 2011; Privitera, Renninger, Carney, Klein, & Aguilar, 2010; Stelmack & Siddle, 1982). Pupil dilation also reliably co-occurs with musical chills (Laeng, Eidet, Sulutvedt, & Panksepp, 2016) – a physiological phenomenon evoked by exposure to emotionally relevant and familiar pieces of music (Harrison & Loui, 2014) and hypothesized to reflect autonomic arousal. Underlying these effects is the increasingly well understood link between non-luminance-mediated change in pupil size and the brain’s neuro-transmitter (specifically ACh and NE; Berridge & Waterhouse, 2003; Joshi, Li, Kalwani, & Gold, 2016; Reimer et al., 2014, 2016; Sara, 2009; Schneider et al., 2016) mediated salience and arousal network.

In particular, abrupt changes in pupil size are commonly observed in response to salient (Liao, Kidani, Yoneya, Kashino, & Furukawa, 2016; Wang & Munoz, 2014) or surprising (Einhäuser, Koch, & Carter, 2010; Lavín, San Martín, & Rosales Jubal, 2014; Preuschoff et al., 2011) events, including in the auditory modality. Work in animal models has established a link between such phasic pupil responses and spiking activity within norepinephrine (alternatively noradrenaline, NE) generating cells in the brainstem nucleus locus coeruleus (LC). The LC projects widely across the brain and spinal cord (Samuels & Szabadi, 2008; Sara & Bouret, 2012) and is hypothesized to play a key role in regulating arousal. Phasic pupil responses are therefore a good measure of the extent to which a stimulus is associated with increased arousal or attentional engagement (Bradley et al., 2008; Partala, Jokiniemi, & Surakka, 2000; Wang, Blohm, Huang, Boehnke, & Munoz, 2017). Here we aim to understand whether and how familiarity drives pupil responses.

Pupil dilations have received much attention in recognition paradigms, analogue to the previously elaborated designs in which subjects are first exposed to a list of stimuli during a learning phase and subsequently asked to identify old items during the recognition stage (Võ et al., 2008). When identifying old words, participants’ pupils tend to dilate more than when confronted with novel ones, a phenomenon which is referred to as the *pupil old/new* effect (Brocher & Graf, 2017; Heaver & Hutton, 2011; Kafkas & Montaldi, 2015). In one of their experiments, Otero et al. (2011) replicated this finding via the use of spoken words, extending this effect onto the auditory domain. The specific timing of these effects is not routinely reported. The bulk of previous work used analyses limited to measuring peak pupil diameter (e.g. Brocher & Graf, 2017; Kafkas & Montaldi, 2011, 2012; Papesh, Goldinger, & Hout, 2012) or average pupil diameter change over the trial interval (Heaver & Hutton, 2011; Otero et al., 2011). Weiss et al. (2016) played a mix of familiar and unfamiliar folk melodies to subjects and demonstrated greater pupil dilations in response to the known-as opposed to the novel stimuli. In this study the effect emerged late, around 6 seconds after stimulus onset. However, this latency may be driven by the characteristics of their stimuli (excerpts ranged in length from 12 to 20 seconds), as well as the fact that melody recognition may take longer than timbre-based recognition.

Here we combine EEG and Pupillometry to investigate the temporal dynamics of the neural processes that underlie the differentiation of familiar from unfamiliar music. In contrast to previous work, which has quantified changes in pupil diameter (the so called ‘pupil dilation response’), we focus on pupil dilation events (see Methods). This approach is substantially more sensitive in the temporal domain and allowed us to tap early activity with the putative salience network. Our experimental paradigm consisted of exposing passively listening participants to randomly presented short snippets from a familiar and unfamiliar song—a design that is reminiscent of the above-mentioned real-world scenarios such as radio channel switching. A control group, unfamiliar with all songs, was also used. We sought to pinpoint the latency at which brain/pupil dynamics dissociate randomly presented familiar from unfamiliar snippets and understand the relationship between brain and pupil measures of this process.

## 2. Methods

### 2.1. Participants

The participant pool encompassed two independent groups: A **main group** (N_main_ = 10; 5 females; Mage= 23.56; SD = 3.71) and a **control group** (N_control_= 12; 9 females; Mage= 23.08; SD = 4.79). All reported no known history of hearing or neurological impairment. Experimental procedures were approved by the research ethics committee of University College London, and written informed consent was obtained from each participant.

### 2.2. Stimuli and procedure

#### 2.2.1. Preparatory stage

Members of the **main group** filled out a music questionnaire, requiring them to list five songs that they have frequently listened to, bear personal meaning and are evocative of positive affect (ratings were expressed on a 5-point scale ranging from ‘1 - Not at all’ to ‘5 - Strong’). One song per subject was then selected and matched with a control song, which was unknown to the participant, yet highly similar in terms of various musical aspects, such as tempo, melody, harmony, vocals and instrumentation. Since we are not aware of an algorithm that matches songs systematically, this process largely relied on the authors’ personal judgments, as well as the use of websites that generate song suggestions (e.g. http://www.spotalike.com, http://www.youtube.com). Importantly, matching was also verified with the control group (see below). Further, we provided each participant with the name as well as a 1500 ms snipped of the matched song. Only if participants were unfamiliar with both, these songs were used as matched ‘unfamiliar’ songs. Upon completion, this procedure resulted in ten dyads (one per participant), each containing one ‘familiar’ and one ‘unfamiliar’ song. See Table 1 for information about the songs selected for the experiment.

**Table 1:**
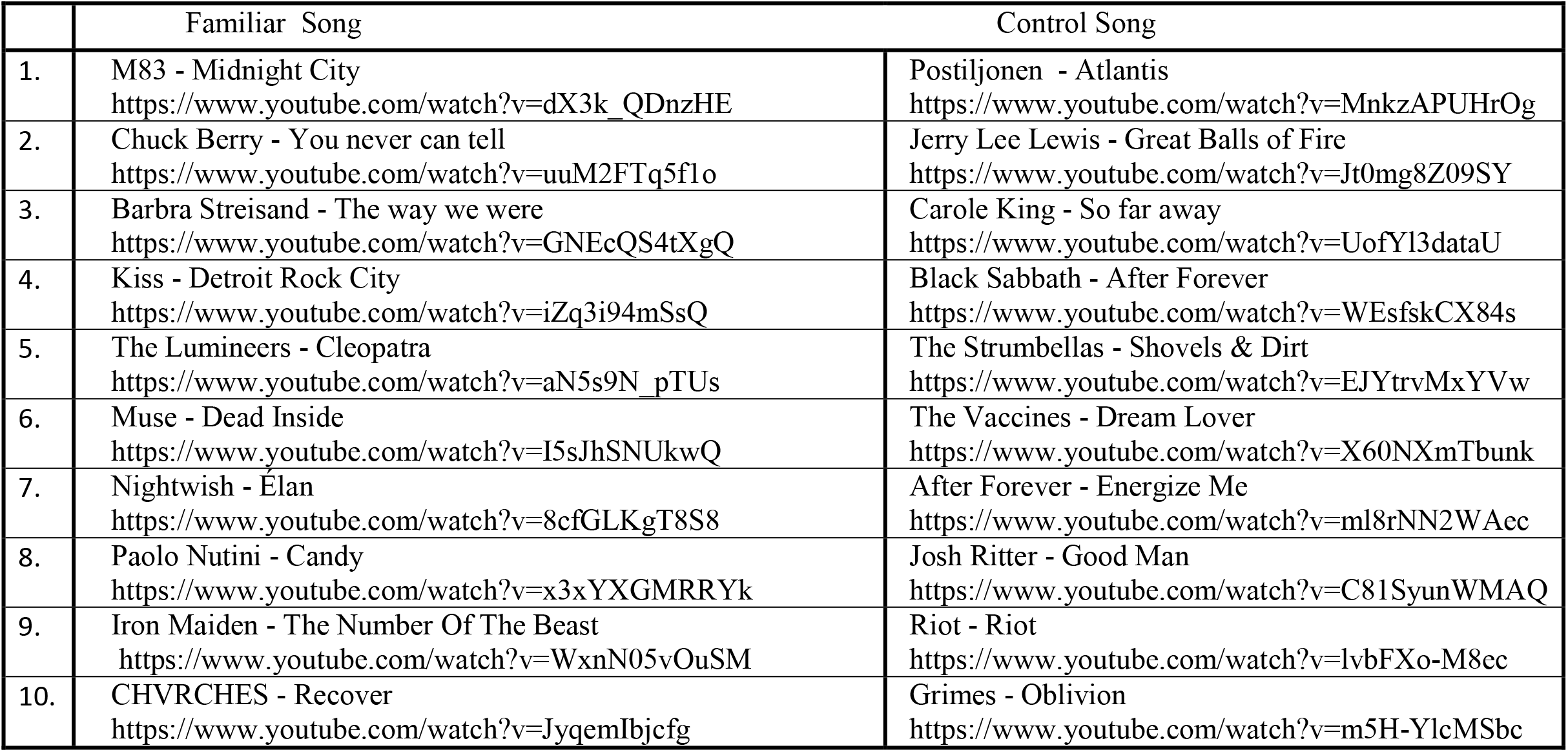
List of song dyads (‘Familiar’ and ‘Unfamiliar’) used in this study. Song were matched for style and timbre quality as described in the methods section. The songs were selected based on input from the ‘main’ group. The ‘control’ group were unfamiliar with all 20 songs.

Participants were selected for the **control group** based on non-familiarity with any of the ten dyads. To check for their eligibility before entering the study, they had the opportunity to inspect the song list as well as to listen to corresponding excerpts (1500 ms). The subjects in the control group (comprised of international students at UCL and hence relatively inexperienced with Western popular music) were unfamiliar with all songs used in this experiment, therefore the distinction between familiar and unfamiliar conditions does not apply to them.

#### 2.2.2. Stimulus generation and testing paradigms

The beginning and end of each song, which typically constitute silent or only gradually rising parts of the instrumentation, were cut out. Both songs from each pair were divided into snippets of 750 ms. Out of these, 100 snippets were randomly chosen for each song. These song snippets were then used in two paradigms: (1) a passive listening task, as well as (2) an active categorization task. In the passive listening task, subjects listened to snippets from each dyad (in separate blocks) in random order whilst their brain activity was recorded with EEG and their pupil diameters with an infrared eye-tracking camera. Each block contained 200 trials, 100 of each song presented in random order with an interstimulus interval (ISI) randomized between 1000 ms and 1500 ms. This resulted in a total duration of 400 seconds per block. Participants from the **main group** were presented with only one block (pertaining to the dyad that contained their familiar song and the matched non-familiar song). Participants from the **control group** listened to all 10 dyads, in random order, each in a separate block.

The active categorization task was also divided into 1 block per dyad. In each block, participants were presented with 20 trials, each containing a random pairing of snippets from each song, separated by 750 ms. They were instructed to indicate whether the two snippets were from the same song or from different ones, by pressing the corresponding buttons on the keyboard (with no time limit imposed). Same as for the passive listening task, subjects from the main group performed only one block, associated with their pairing of ‘familiar/unfamiliar’ songs. Subjects from the control group completed 10 blocks in random order.

#### 2.2.3. Procedure

Participants were seated, with their heads fixed on a chinrest, in a dimly lit and acoustically shielded testing room (IAC Acoustics, Hampshire, UK). They were distanced 61 cm away from the monitor and 54 cm away from two loudspeakers, arranged at an angle of 30° to the left and right of the subject. Subjects were instructed to continuously fixate on a white cross on grey background, presented at the center of a24-inch monitor (BENQ XL2420T) with resolution of 1920×1080 pixels and a refresh rate of 60 Hz. They first engaged in the passive listening task followed by the active categorization task. For both tasks, subjects were instructed to attentively listen to the snippets. EEG and eye-tracking were recorded during the first task, but not during the second, which was of purely behavioral nature.

### 2.3. EEG acquisition, preprocessing and analysis

EEG recordings were obtained using a Biosemi Active Two head-cap 10/20 system with 128 scalp channels. Eye movements and blinks were monitored using 2 additional electrodes, placed on the outer canthus and infraorbital margin of the right eye. The data were recorded reference-free with a passband of 0.016-250 Hz and a sampling rate of 2048 Hz. After acquisition, pre-processing was done in Matlab (The MathWorks Inc., Natick, MA, USA), with EEGLAB (http://www.sccn.ucsd.edu/eeglab/; Delorme & Makeig, 2004) and FieldTrip software (http://www.ru.nl/fcdonders/fieldtrip/; Oostenveld et al., 2010). The data were downsampled to 128 Hz, low pass filtered at 40 Hz and re-referenced to the average across all electrodes. The data were not high pass filtered, to preserve low frequency activity (Kappenman & Luck, 2012), which is relevant when analyzing sustained responses. The data were segmented into stimulus time-locked epochs ranging from −500 ms to 1500 ms. Epochs containing artefacts were removed on the basis of summary statistics (variance, range, maximum absolute value, z-score, maximum z-score, kurtosis) using the visual artefact rejection tool implemented in Fieldtrip. On average, 2.1 epochs per song pair in the main group and 4.9 epochs in the control group were removed. Artefacts related to eye movements, blinks and heartbeat were identified and removed using independent component analysis. Subsequently the data were re-referenced to the mean of all channels, averaged over epochs of the same condition and baseline-corrected (200 ms preceding stimulus onset).

A cluster-based permutation analysis which takes spatial and temporal adjacency into account (Maris & Oostenveld, 2007; Oostenveld et al., 2010) was used to investigate potential differences between the EEG responses to ‘familiar’ and ‘unfamiliar’ snippets within controls and main participants over the entire epoch length. The significance threshold was chosen to control family-wise error-rate (FWER) at 5%.

### 2.4. Pupil measurement and analysis

Gaze position and pupil diameter were continuously recorded by an infrared eye-tracking camera (Eyelink 1000 Desktop Mount, SR Research Ltd.), positioned just below the monitor and focusing binocularly at a sampling rate of 1000 Hz. The standard five-point calibration procedure for the Eyelink system was conducted prior to each experimental block. Due to a technical fault that caused missing data, seven control participants were excluded from the pupillometry analysis, leaving five valid participants (5 females, Mage = 23.21, SD = 4.37) in the control group. No participant was excluded in the main group.

The standard approach for analyzing pupillary responses involves across trial averaging of pupil diameter as a function of time. This is usually associated with relatively slow dynamics (Hoeks & Levelt, 1993; Liao, Yoneya, Kidani, Kashino, & Furukawa, 2016; Murphy, Robertson, Balsters, & O’connell, 2011; Wang & Munoz, 2015) which are not optimal for capturing potentially rapid effects within a fast-paced stimulus. Instead, the present analysis focused on examining pupil event rate. This analysis captures the incidence of pupil dilation events (Joshi et al, 2016), irrespective of their amplitude, and therefore provides a sensitive measure of subtle changes in pupil dynamics that may be evoked by the familiar vs. non-familiar stimuli. Pupil dilation events were extracted from the continuous data by identifying the instantaneous positive sign-change of the pupil diameter derivative (i.e. the time points where pupil diameter begins to positively increase). To compute the incidence rate of pupil dilation events, the extracted events were convolved with an impulse function (see also Joshi, Li, Kalwani, & Gold, 2016; Rolfs, Kliegl, & Engbert, 2008), paralleling a similar technique for computing neural firing rates from neuronal spike trains (Dayan & Abbott, 2002). For each condition, in each participant and trial, the event time series were summed and normalized by the number of trials and the sampling rate. Then, a causal smoothing kernel ω(τ)=α^2×τ×e^(-ατ) was applied with a decay parameter of α = 1/50 ms (Dayan & Abbott, 2002; Rolfs et al., 2008; Widmann, Schröger, & Wetzel, 2018). The resulting time series was then baseline corrected over the pre-onset interval. For each condition, the pupil dilation rate averaged across participants is reported here.

To identify time intervals in which the pupil dilation rate was significantly different between the two conditions, a nonparametric bootstrap-based statistical analysis was used (Bradley & Tibshirani, 1994): For the main group, the difference time series between the conditions was computed for each participant, and these time series were subjected to bootstrap re-sampling (1000 iterations). At each time point, differences were deemed significant if the proportion of bootstrap iterations that fell above or below zero was more than 99% (i.e. p<0.01).

Two control analyses were also conducted to verify the effects found in the main group. Firstly, **permutation analysis on the data from the main group**: in each iteration (1000 overall), 10 participants were selected with replacement. For each participant all trials across conditions were randomly mixed and artificially assigned to the ‘familiar’ or the ‘unfamiliar’ condition (note that theses labels are meaningless in this instance). This analysis yielded no significant difference between conditions. A second control analysis examined **pupil dynamics in the control group**. Data for each control participant consisted of 10 blocks (one per dyad), and these were considered as independent data sets for this analysis, resulting in 50 control datasets. To calculate the differences between conditions, 10 control datasets were selected with replacement from the pool of 50 and used to compute the mean difference between the two conditions. From here, the analysis was identical to the one described for the main group. This analysis also yielded no significant difference between conditions.

## 3. Results

### 3.1. Music questionnaire

Subjects in the **main group** were highly familiar with the selected songs (Familiarity Score; M = 3.91; SD = 0.85) and reported increased emotional ratings for items such as ‘excited’ (M = 3.50; SD = 1.08), ‘happy’ (M = 3.60; SD = 1.26), and ‘memory inducing’. Conversely, control subjects gave ratings that were indicative of low familiarity for all the songs (M = 1.05; SD = 0.09), as well as their respective artists in general (M = 1.10; SD = 0.14).

### 3.2. EEG

The general EEG response to the sound snippets (collapsed across all participants and conditions) is shown in Figure 1. The snippets evoked a characteristic onset response, followed by a sustained response. The onset response was dominated by P1 (at 71 ms) and P2 (at 187 ms) peaks, as is commonly observed for wide-band signals (e.g. Chait et al, 2004).

**Figure 1.**
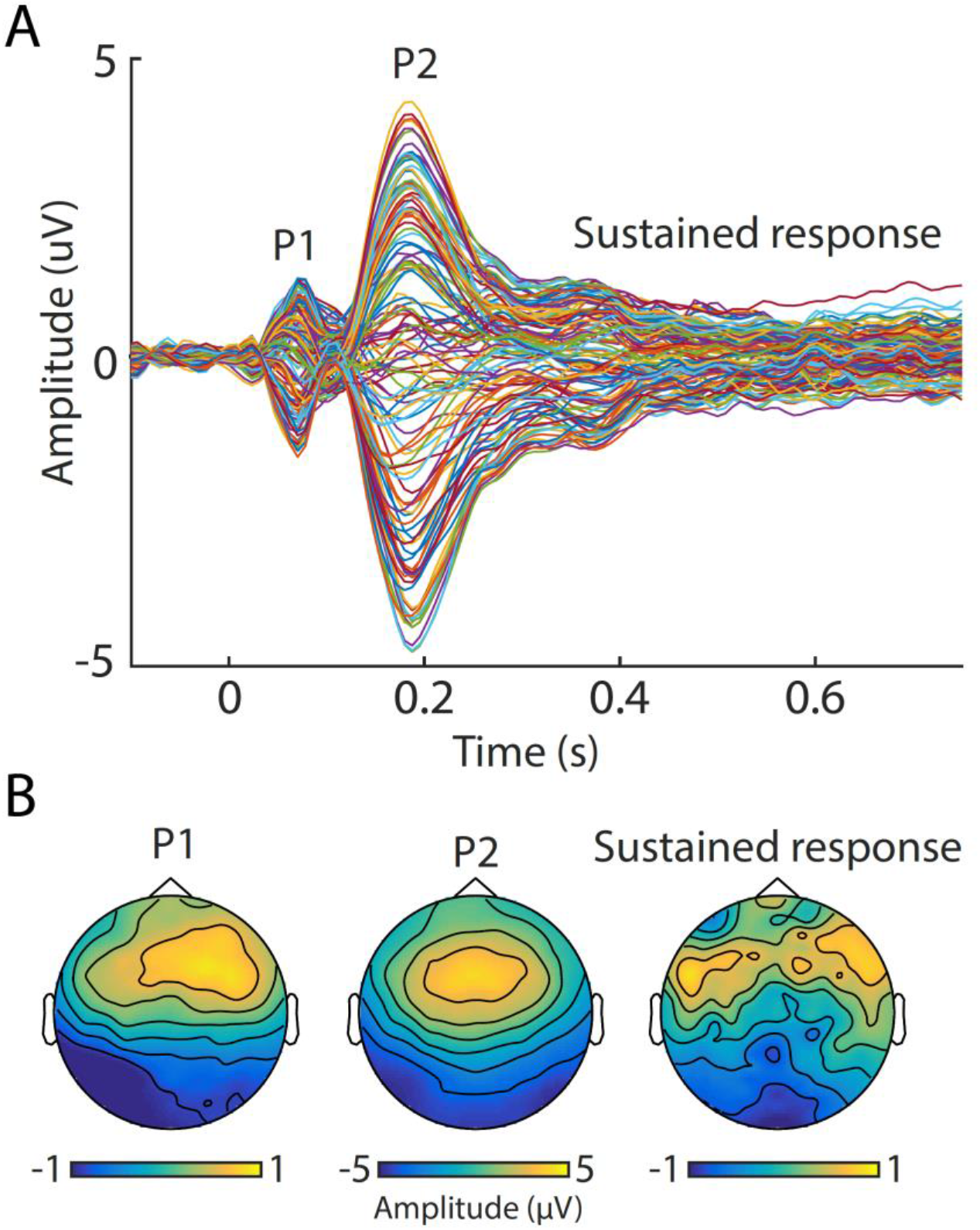
Grand-average event-related potential, averaged across the main and control group, as well as familiar and unfamiliar songs, demonstrating the brain response to the music snippets. (A) Time-domain representation. Each line represents one EEG channel. (B) Topographical representation of the P1 (71 ms), P2 (187 ms), as well as the sustained response (plotted is average topography between 300-750ms).

### 3.3.1. Control group

The main purpose of the control group was to verify that any significant differences which are potentially established for main participants were due to our familiarity manipulation and not caused by any acoustic differences between the songs in each dyad. Because the control group participants were unfamiliar with the songs, we expected no differences in brain activity. However, the cluster-based permutation test revealed significant clusters in dyad 2 (mean t = 3.15) and 5 (mean t = 2.84). Those dyads were excluded from the subsequent main group analysis.

### 3.3.2. Main group

Comparing responses to ‘familiar’ and ‘unfamiliar’ snippets within the main group, we identified two clusters of channels showing a significant difference between the two conditions. A centrally located cluster of 26 channels showing a significant difference between conditions from 540 to 750 ms (mean t = −4.52), and a right fronto-temporal cluster of 20 channels, showing significant difference between 350 to 750 ms (mean t =4.15). In both clusters, responses to unfamiliar snippets evoked stronger activity (more negative in the first cluster and more positive in the second cluster) than familiar snippets (Figure 2).

**Figure 2.**
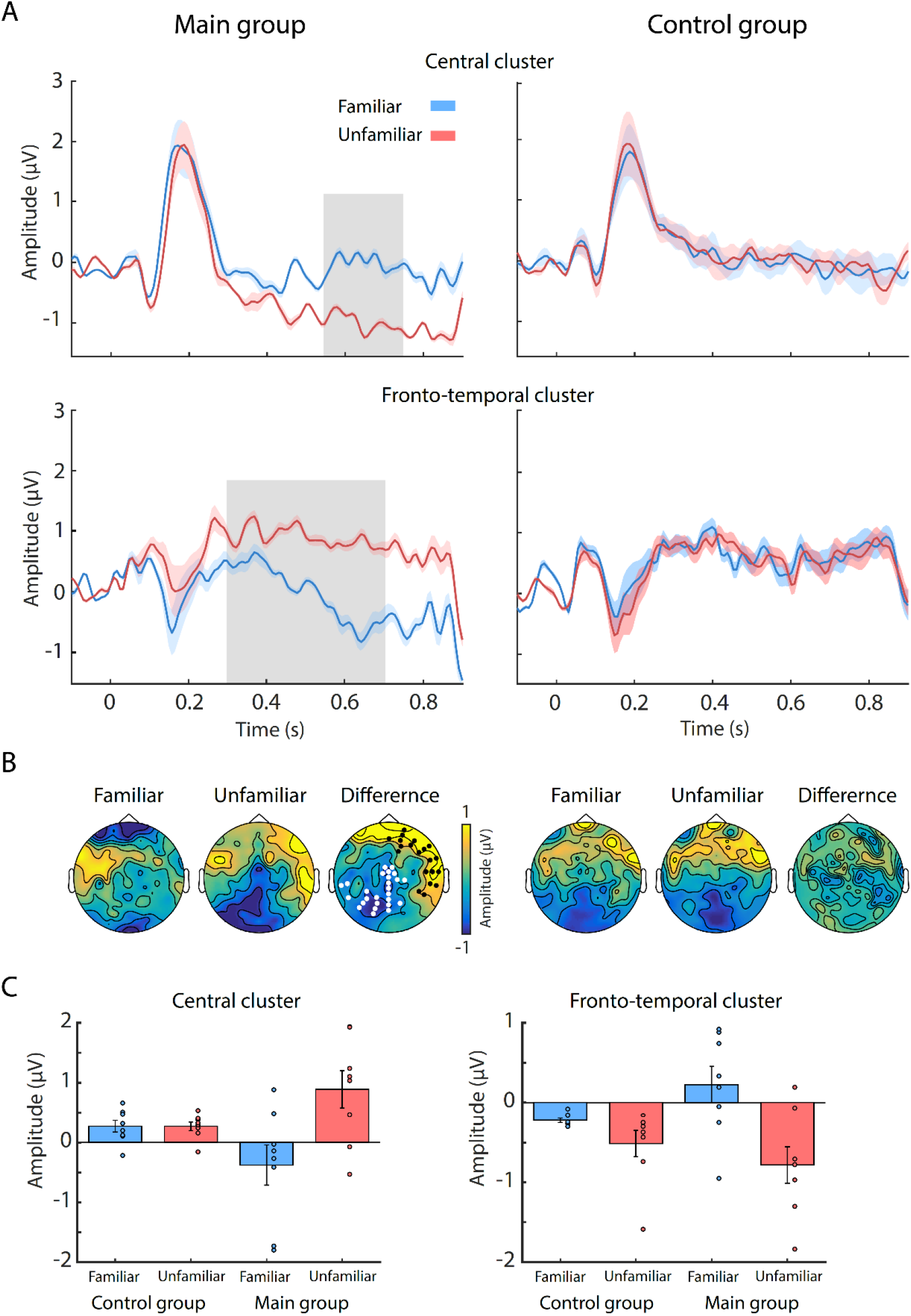
Event-related potential results - differences between ‘familiar’ and ‘unfamiliar’ snippets in the main, but not control, group. (A) Time-domain ERPs for the central cluster (top row) and fronto-central cluster (bottom row), separately for the main (left column) and control (right column) group. Solid lines represent mean data for familiar (blue) and unfamiliar (red) songs (note that this labeling only applies to the ‘main’ group; both songs were unfamiliar to the control listeners). Shaded areas represent standard error of the mean. Significant differences between conditions, as obtained via cluster-based permutation tests, are indicated by grey boxes. (B) Topographical maps of the familiar and unfamiliar ERP responses as well as their difference (from 350 to 750 ms), separately for the main (left column) and control (right column) group. White and black dots indicate electrodes belonging to the central and fronto-temporal cluster, respectively. (C) Barplots of mean ERP amplitudes, as entered into the ANOVA. Main group, but not control group, showed significantly larger responses to unfamiliar song snippets, at both the central and the fronto-temporal cluster. Errorbars represent standard error of the mean, coloured dots represent individual subjects’ mean.

To confirm that this effect is specific to the main group, the identified clusters were used as ROIs for a factorial ANOVA across groups, with familiarity (familiar/unfamiliar) as independent within-subjects factor and the average EEG response amplitudes within the cluster as dependent variable. The interaction was significant for both the central cluster (F (1, 14) = 73.56, p < .001) as well as the fronto-temporal cluster (F (1, 14) = 37.91, p < .001) and. More precisely, compared to controls, the main group showed increased negativity in the central regions and increased positivity in a right fronto-temporal area in response to unfamiliar snippets (Figure 2C).

### 3.3. Pupil dilation

#### 3.3.1. Control group

Figure 3A (bottom) shows the pupil dilation rates for familiar and unfamiliar snippets in the control group. In response to the auditory stimuli, the pupil dilation rate increased shortly after the onset of a snippet, peaking at around 400ms, before returning to baseline around 550ms post-onset. No difference was observed between the two conditions throughout the entire epoch.

**Figure 3.**
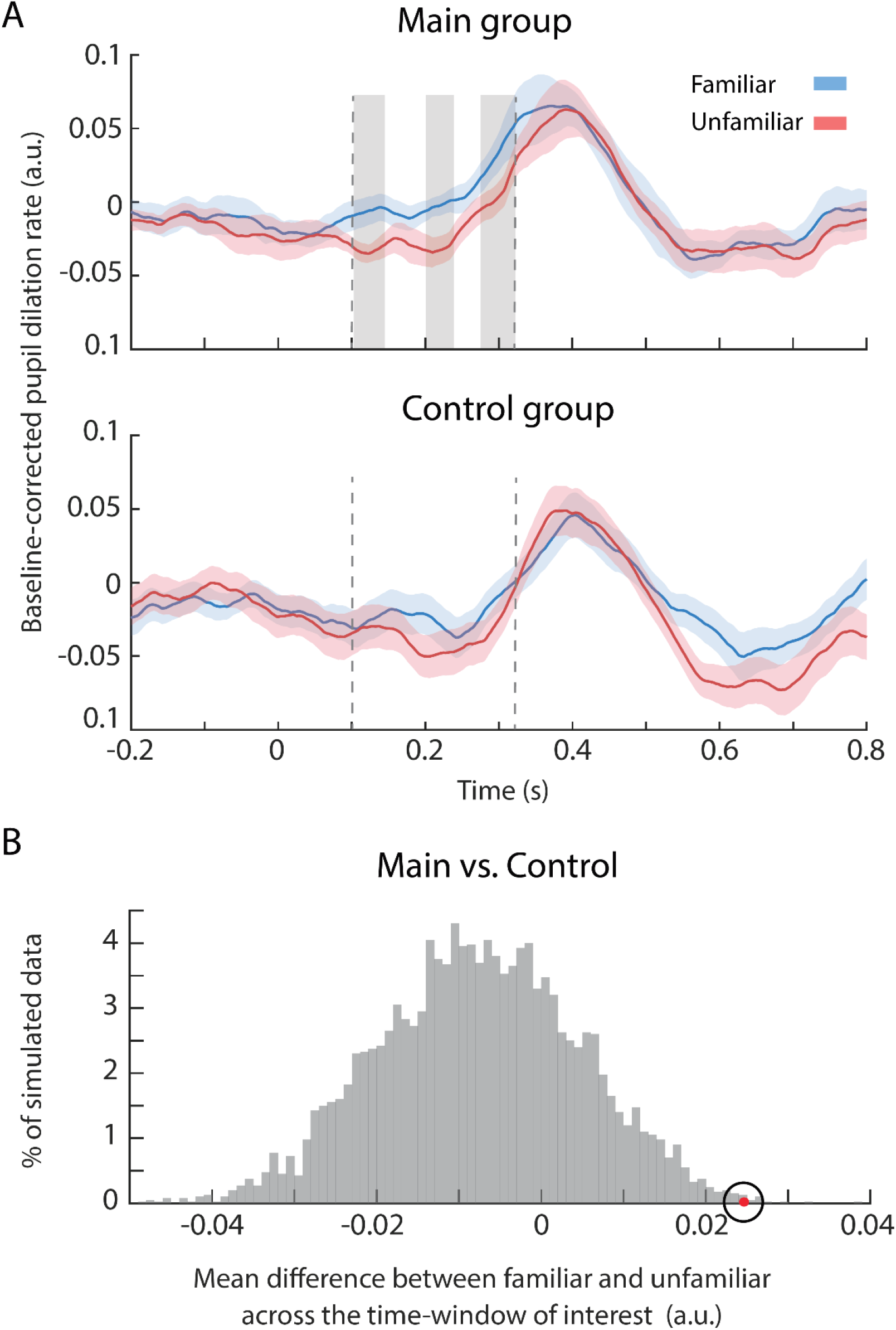
Pupil dilation rate to familiar and unfamiliar snippets. (A) Top: Main group. The solid curves plot the average pupil dilation rate across subjects for familiar (blue) and unfamiliar (red) conditions. The shaded area represents one standard deviation of the bootstrap resampling. The grey boxes indicate time intervals where the two conditions were significantly different (114–137, 217–230, and 287–314ms). The dashed lines indicate the time interval use for the resampling statistic in B. Bottom: Control group. The solid curves plot the average PD rate over 10 randomly selected control datasets. The shaded area represents one standard deviation of the bootstrap resampling. No significant differences were observed (B) Results of the resampling test to compare the difference between familiar and unfamiliar conditions (during the time interval 108-315 ms, indicated via dashed lines in A) across the main and control groups. The grey histogram shows the distribution of differences between conditions for the control group. The red dot indicates the observed difference for the main group.

#### 3.3.2. Main group

When compared with unfamiliar conditions, familiar snippets were associated with a higher pupil dilation rate from 108–315ms post sound onset (Figure 3A, top). This significant interval was absent in the shuffled data (see methods).

We also directly compared the difference between ‘familiar’ and ‘unfamiliar’ conditions between the two groups during the time interval (108- 315 ms) identified as significant in the main group analysis. This was achieved by computing a distribution of differences between conditions based on the control group data (H0 distribution). On each iteration (5000 overall) 10 datasets were randomly drawn from the control pool and used to compute the difference between conditions during the above interval. The grey histogram in Figure 3B, shows the distribution of these values. The mean difference from the main group, indicated via the red dot, lies well beyond this distribution (p=0.0022), confirming that the effect observed for the main group was different from that in the control group.

### 3.4. Behavioral task

The behavioral task aimed to verify whether subjects were able to differentiate between familiar and unfamiliar snippets, and whether participants in the **main group** (who were highly familiar with one song in a pair) performed better than controls.

Main subjects correctly identified whether or not the two presented snippets were from the same song in 92% of trials, whereas controls in 79% of trials. One-sample t-tests revealed that these scores are at above-chance levels, t(9) = 13.61, p< .00001 for controls, and t(9) = 18.11, p< .000001 for the main group. An independent samples t-test revealed that main subjects scored significantly higher than controls t(18) = 6.19, p< .00001. Therefore, the behavioral results suggest that both groups were able to differentiate the songs in each dyad well above chance. That the control group participants achieved above chance performance suggests that there was sufficient acoustic information in the snippet to distinguish the songs.

## 4. Discussion

We used EEG and eye tracking to identify brain responses which distinguish between familiar and unfamiliar music. To tap rapid recognition processes, matched ‘familiar’ and ‘unfamiliar’ songs were divided into brief (750 ms) snippets which were presented in a mixed, random, order to passively listening subjects. We demonstrate that despite the random presentation order, pupil and brain responses swiftly distinguished between snippets taken from familiar vs. unfamiliar songs, suggesting rapid underlying recognition mechanisms. Specifically, we report two main observations: (1) pupil responses showed greater dilation rate to snippets of familiar music from ~100-300 ms post stimulus (2) brain activity measured with EEG showed a differentiation between responses to familiar and unfamiliar music snippets from 350 ms post onset, but, notably, in the opposite direction from that observed with pupillometry: Unfamiliar music snippets evoked stronger responses.

The implications of these results for our understanding of how music experiences are coded in the brain (Koelsch, 2018) are discussed below. But to start with we outline several important limitations which the reader must keep in mind: Firstly,’familiarity’ is a multifaceted concept. In the present study, songs were explicitly selected to evoke positive feelings and memories. Therefore, for the ‘main’ group the ‘familiar’ and ‘unfamiliar’ songs did not just differ in terms recognizability but also in terms of emotional engagement and affect. Whilst we continue to refer to the songs as ‘familiar’ and ‘unfamiliar’, the effects we observed may also be linked with these other factors. Secondly, though we took great care in the song matching process, ultimately this was done by hand due to lack of availability of appropriate technology. Advancements in automatic processing of music may improve matching in the future. Alternatively, it may be possible to work with a composer to ‘commission’ a match to a familiar song. Another major limitation is the fact that it was inevitable that participants in the main group were aware of the aim of the study and might have listened with an intent that is different from that in the control group. This limitation is hard to overcome, and the results must be interpreted in this light.

Pupil responses revealed a very early effect, dissociating the ‘familiar’ and ‘unfamiliar’ snippets from 107ms after onset. To our knowledge, this is the earliest pupil old/new effect documented, though the timing is broadly consistent with previous behaviorally derived estimates, which place minimum identification time for music at 100-250 ms (Schellenberg et al. 1999; Bigand, 2009). This rapid recognition likely stems from remarkable sensitivity to, and memory of, the timbre of the familiar songs (Suied et al., 2014; Agus et al, 2012).

Research in animal models has linked phasic pupil dilation events with increased firing in the LC (Joshi et al., 2016). The present results can therefore be taken to indicate that the LC was differentially activated as early as 100 ms after sound onset, possibly through projections from the inferior colliculus (where timbre cues may be processed; Zheng & Escabi, 2008, 2013) or other subcortical structures such as the amygdala, which is known to activate the LC (Bouret, Duvel, Onat, & Sara, 2003). The Amygdala has been extensively implicated in music processing, especially for familiar pleasure-inducing music such as the one used here (Blood & Zatorre, 2001; Salimpoor et al., 2013).

It is notable that the pupil effects were restricted to the early portion of the trial. A possible explanation is that later in the trial, the relatively subtle effects of familiarity were masked by the more powerful effects associated with processing the snippet as a whole.

In contrast to the pupil-based effects, in the EEG analysis familiarity was associated with *reduced* responses between 350 to 750 ms. This finding also conflicts with previous demonstrations of increased EEG activation to familiar items -the so called old/ new effect (Kappenman & Luck, 2012; Wagner et al., 2005). The discrepancy with the previous literture might be due to the fact that here subjects were listening passively and not making active recognition judgments, as it was the case in the majority of past paradigms (e.g. Sanquist et al., 1980). Further, whilst previously reported ‘old/new’ effects occurred post stimulation, here the difference manifested during ongoing listening.

The present study does not have sufficient data for reliable source analysis, however from the overall field maps (Figure 2B) it appears that the pattern of increased activation for ‘unfamiliar’ snippets encompasses areas such as the right superior temporal gyrus (rSTG), right inferior and middle frontal gyri (rIFG/rMFG), which have been implicated in the recognition of familiarity notably in the context of voices (Blank, Wieland, & von Kriegstein, 2014; von Kriegstein, Eger, Kleinschmidt, & Giraud, 2003) and are therefore potential neural generators of the present effect. Indeed, Zäske et al. (2017) demonstrated that exposure to unfamiliar voices entailed an increased activation in those areas. These increases in activation may be associated with explicit memory-driven novelty detection or else reflect more general increase in activation related to attentional capture, or listening effort associated with processing of unfamiliar sounds. Both the rIFG and rMFG have been implicated in a network that allocates processing resources to external stimuli of high salience (Corbetta, Patel, & Shulman, 2008; Corbetta & Shulman, 2002; Shulman et al., 2009).

Notably, no differences were observed in the control group, despite the behavioral evidence that indicated that they can differentiate the ‘familiar’ and ‘unfamiliar’ songs. This suggests that whilst there may have been enough information for active perceptual scanning to indicate acoustic or stylistic differences between songs which would lead to above chance recognition, these processes did not affect the presently observed brain responses during passive listening.

Together, the data reveal early effects of familiarity in the pupil dynamics measure and later (opposite direction) effects in the EEG brain responses. The lack of earlier effects in EEG may result from various factors, including that early brain activity may not have been measurable with the current setup. This could happen if effects are small, or sources not optimally oriented for capture by EEG. In particular, as discussed above, the rapid pupillometry effects are likely to arise from sub-cortical recognition pathways and are therefore not measurable on the scalp. Future research combining sensitive pupil and brain imaging measurement is required to understand the underlying network. One possible hypothesis, consistent with the present pattern of results, is that familiar snippets are recognized rapidly, likely based on timbre cues, mediated by fast acting sub-cortical circuitry. This rapid dissociation between ‘familiar’ and ‘unfamiliar’ snippets leads to later increased processing associated with the novel input e.g. as expected by predictive coding views of brain function (de Lange, Heilbron, & Kok, 2018; Friston & Kiebel, 2009; Heilbron & Chait, 2018) whereby surprising, unknown stimuli require more processing than familiar, expected, signals.

## Acknowledgements

This research was supported by a EC Horizon 2020 grant and a BBSRC international partnering award to MC.

## Competing Interests

The authors declare no competing interests.

## Author Contributions

RJ: designed the study, conducted the research; analyzed the data; wrote the paper

UP: designed the study, conducted the research; analyzed the data; wrote the paper

MY: conducted pilot experiments; commented on draft

SZ: analyzed the data; wrote the paper

MC: supervised the research; obtained funding; designed the study; wrote the paper

## Data Accessibility

Upon acceptance the (de-identified) data supporting the results in this manuscript (together with any relevant analysis scripts) will be archived in a suitable online depository.

